# The shifting of dominating roles between structural cells and immune cells are key regulators of human adipose tissues aging

**DOI:** 10.1101/2021.02.15.431264

**Authors:** Wenyan Zhou, Junxin Lin, Xueqing Hu, Xudong Yao, Hongwei Ouyang

## Abstract

Adipose tissue is a highly dynamic organ with complex cellular composition. Aging induces adipose tissue function decline and relocation of peripheral adipose tissue to abdominal compartment, which often associated with inflammation and metabolic disorders. Here we performed single-cell RNA sequencing to comprehensively and unbiasedly deconvolve how subcutaneous adipose tissue (SAT) responses to aging. We collected >25,000 stromal vascular cells from abdominal and gluteofemoral SAT of young and old donors. Analyses of transcription signatures and cell networks uncovered impaired adipogenesis and extracellular matrix synthesis capacity of APC, altered metabolic phenotype of immune cells and shifted tissue-dominating cells that can be used to predict adipose tissue aging. We also reported aging-associated distinct transcriptional program between gluteofemoral SAT and abdominal SAT. Our work thus reveals unanticipated cellular, immunological, metabolic and site-specific aspects of human adipose tissues aging process, providing valuable resource for better understanding of aging-associated adipose tissue dysfunction.

## Introduction

Adipose tissue distributes throughout the body. It is an extremely complex organ with profound effects on physiology and pathophysiology of human body, and plays critical roles in mechanical support, temperature regulation, metabolic and immune homeostasis ^1^. Aging is increasingly recognized as a kind of systemic chronic disease, leading to the functional decline of vital tissues. Age-dependent adipose tissue dysfunction is associated with metabolic declines, heterotopic fat accumulation and chronic systemic inflammation, increasing the risk of diseases like diabetes, cardiovascular disease, and even cancer ^2^.

Aging changes the mass and distribution of adipose tissue throughout the body. Body weight increases with age and fat mass peaks occur in middle or early old age ^3-5^. With increasing age, peripheral subcutaneous adipose tissue tends to be lost, while visceral adipose tissue (VAT) tends to be preserved ^6,7^. Similar to the VAT, the fat mass of the abdominal subcutaneous adipose tissue (ASAT) increases with advancing age in both male and female ^8,9^, which positively correlates with cardiovascular disease and type 2 diabetes mellitus ^10,11^. Recent studies have shown that adipose progenitor differentiation, adipose tissue immune and metabolic tenors are tightly regulated by the crosstalk between adipose progenitor cells and non-adipocyte fraction of human adipose tissue, including immune cells and structural cells ^12,13^. However, how aging influences cell subpopulations and their crosstalk in human adipose tissue across depots is still unclear.

In this study, we present high-throughput single-cell transcriptome analyses of young and old human ASAT and GSAT. Our results reveal the aging-dependent alterations in cell subpopulations, cell-cell interactions, as well as site-specific response to aging of human SAT.

## Results

### Single-cell atlas of human ASAT

We performed single-cell RNA sequencing on stromal vascular fraction (SVF) cells of ASAT from young and aged participants (Figure. 1A). In total, 14,073 single cells derived from the ASAT of 3 young (26.33±3.79 years old) and 3 old (74.33±3.51 years old) donors were analyzed (Supplementary Figure. 1), unsupervised clustering of the gene expression profiles identified 11 cell types, each containing cells from both young and old samples (Figure. 1B, Supplementary Figure. 2). A list of differentially expressed genes (DEGs) that define the clusters are presented in Supplementary Table 1. Analysis of DEGs identified six major clusters of adipose progenitor cells (APC), three clusters of immune cells (IC), a population of vascular endothelial cells (VEC) and a small population of smooth muscle cells (SMC) (Figure. 1B,C).

**Figure 1.**
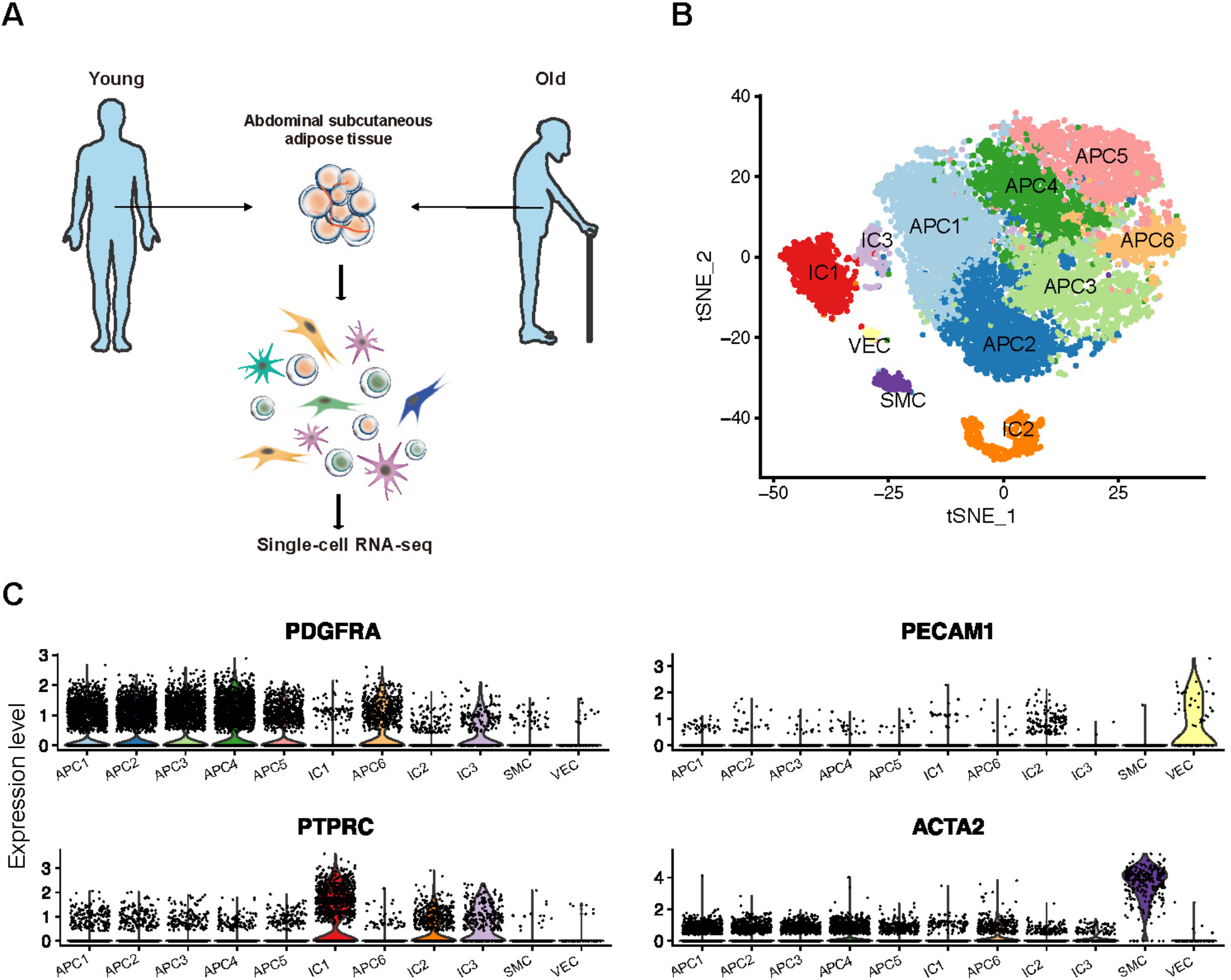
Single-cell RNA-sequencing of human ASAT. (A) Schematic diagram of the experimental workflow. (B) The t-Distributed stochastic neighbor (t-SNE) plot shows unsupervised clustering of 14,073 single-cell transcriptomes. (C) Violin plots show the expression levels of representative cell-type-specific marker genes across all 11 cell types.

**Figure 2.**
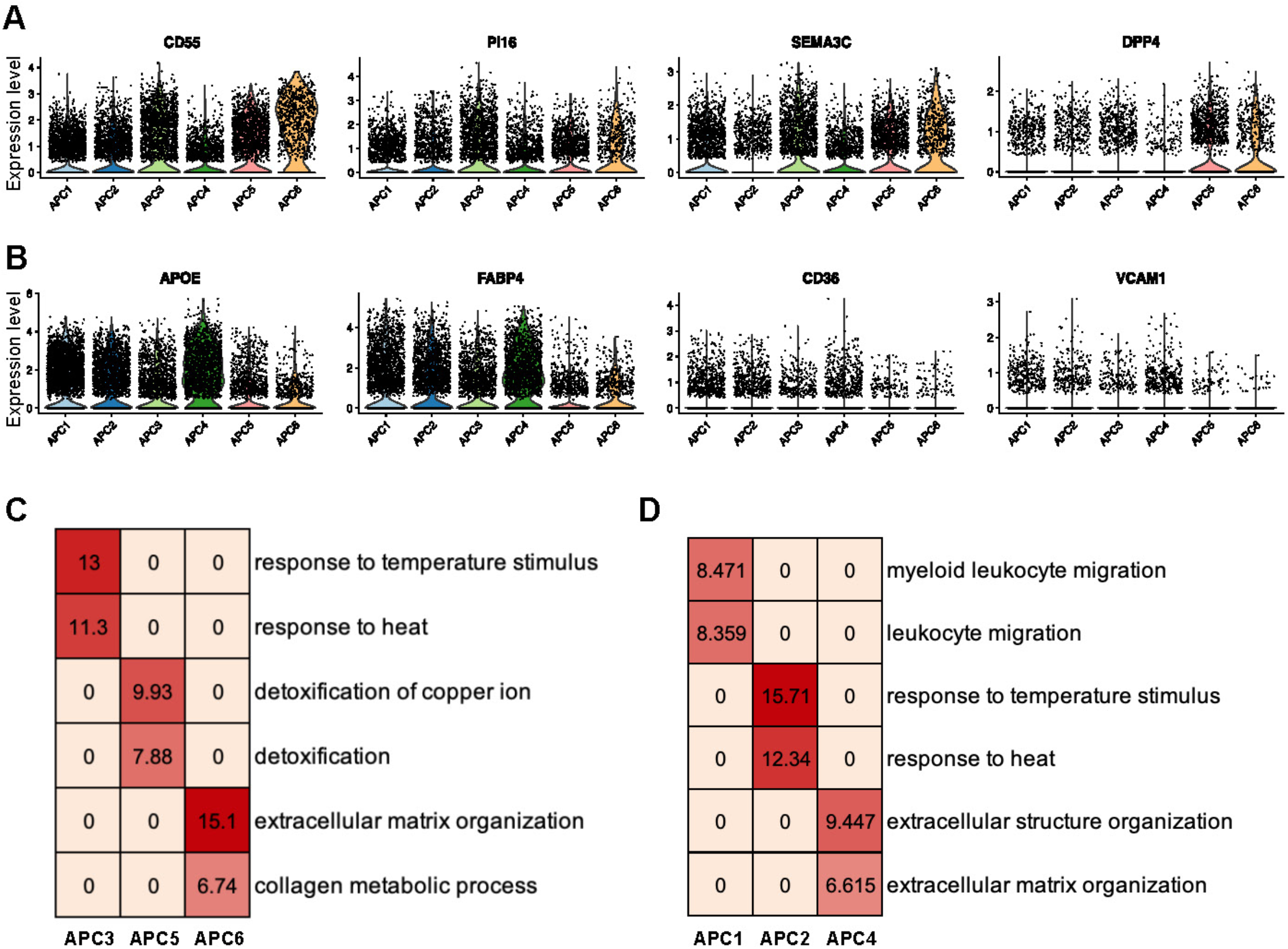
APC heterogeneity of human ASAT. (A) Violin plots show the expression levels of stem cell markers and (B) early adipogenic markers of human APC. (C) Heatmap shows the gene ontology term of differentially expressed genes of stem-like populations APC3, APC5, APC6 and (D) committed preadipocyte populations APC1, APC2, APC4.

Our data showed that APC3, APC5, APC6 expressed higher level of stem cell markers (*CD55, PI16, SEMA3C, DPP4*) (Figure. 2A), while APC1, APC2 and APC4 expressed higher level of early adipogenic markers (*APOE, FABP4, CD36, VCAM1*) ^14-16^ (Figure. 2B). The three stem-like APC clusters could be distinguished by high expression of genes corresponding to temperature response in APC3, detoxification in APC5, and extracellular matrix (ECM) organization in APC6 (Figure. 2C). Within the three committed preadipocyte population, APC2 and APC4 exhibited the similar phenotype with stem-like population APC3 and APC6 respectively, which represents the temperature-effector differentiation trajectory (APC3-APC2) and ECM-effector differentiation trajectory (APC6-APC4). APC1 was featured by high expression of genes associated with leukocyte migration (Figure. 2D). These results reveal the coordination of cell type and maturation stage within human APC populations.

It was suggested that aging increased cellular transcriptional instability, which led to cell fate drift and function loss ^17^. Therefore, we analyzed transcriptional noise following previous work, and identified that transcriptional noise was increased with aging in major cell populations inhabited in human ASAT (Supplementary Figure. 3), which was consistent with the previous findings in mice ^17,18^.

**Figure 3.**
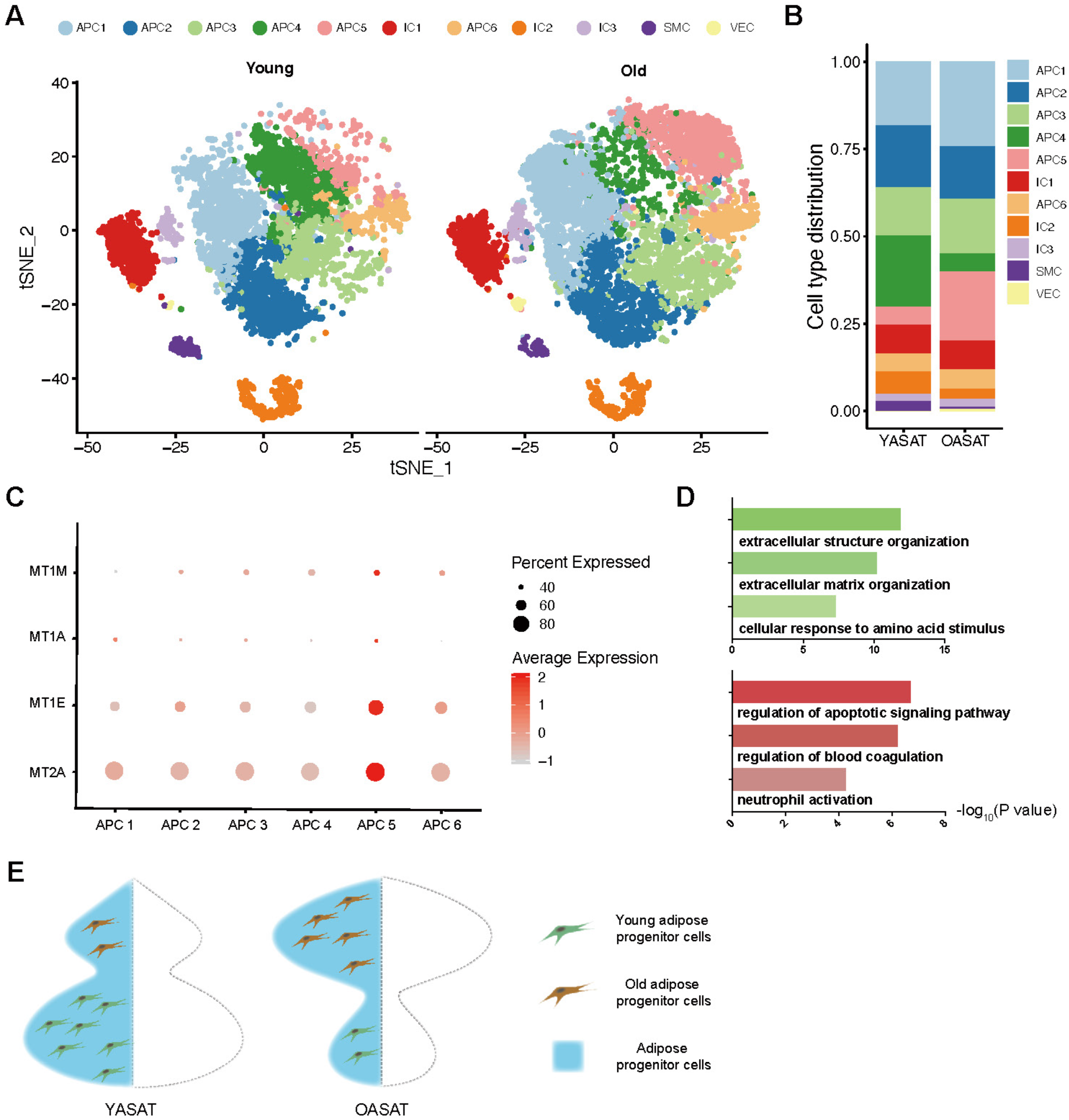
Alterations of APC subpopulations during ASAT aging. (A) The t-SNE plots of cell clusters in young and old ASAT split from Figure.1 B. (B) Cell type distribution of young and old ASAT. (C) Dot plot of the expression of metallothionein genes across APC populations. (D) Representative GO terms of common downregulated (upper plot) and upregulated (lower plot) genes during aging of APC1, APC2, APC3, and APC6. YASAT: young abdominal subcutaneous adipose tissue; OASAT: old abdominal subcutaneous adipose tissue. (E) A schematic representation of the changes of adipose progenitor cell subpopulations in young and old ASAT.

### Aging causes the accumulation of a dysfunctional APC subpopulation

Segregation of the aggregated t-SNE plot of APC demonstrated that committed preadipocyte APC4 was mainly composed of cells from the young ASAT (YASAT), whereas stem-like population APC5 included cells mostly from the old ASAT (OASAT) (Figure. 3A,B). Besides the stem cell property, APC5 expressed high level of metallothionein genes (*MT1A, MT1E, MT1M, MT2A*), which are associated with cell dysfunction ^19^(Figure. 3C). These alterations indicate that aging impeded the differentiation of ECM-trajectory (APC6-APC4), result in the accumulation of dysfunctional stem-like APC5. Although the changes of cellular quantity were minor, the split t-SNE plot further indicated that aging shifted the expression profile of APC1, APC2, APC3, and APC6.

To characterize the common features caused by aging, we analyzed differentially expressed genes for APC1, APC2, APC3, and APC6 across young and aged ASAT, and identified a series of genes commonly changed by aging in these clusters. Only those genes consistently changed (up-regulated or down-regulated) across all these four clusters were defined as common DEGs (Supplementary Table 2). Gene ontology (GO) analysis of the common DEGs of these four APC populations during aging showed that genes related to extracellular structure organization were downregulated, while genes involved in apoptotic signaling pathway, blood coagulation, and neutrophil activation were upregulated (Figure. 3D). These results show that aging induced the expansion of dysfunctional aged adipose progenitor cells and partly impedes the adipogenic differentiation (Figure. 3E), but the ECM synthesis ability was broadly hindered across all APC populations.

### Aging alters the immune and metabolic phenotype of human ASAT

Previous studies have highlighted immune cells alteration responding to adipose tissue aging. To learn more details about immune cell profile of ASAT, we re-clustered cells of IC1-IC3 and identified 8 specific immune cell subpopulations, labelled as ICS1-ICS8 (Figure. 4A).

**Figure 4.**
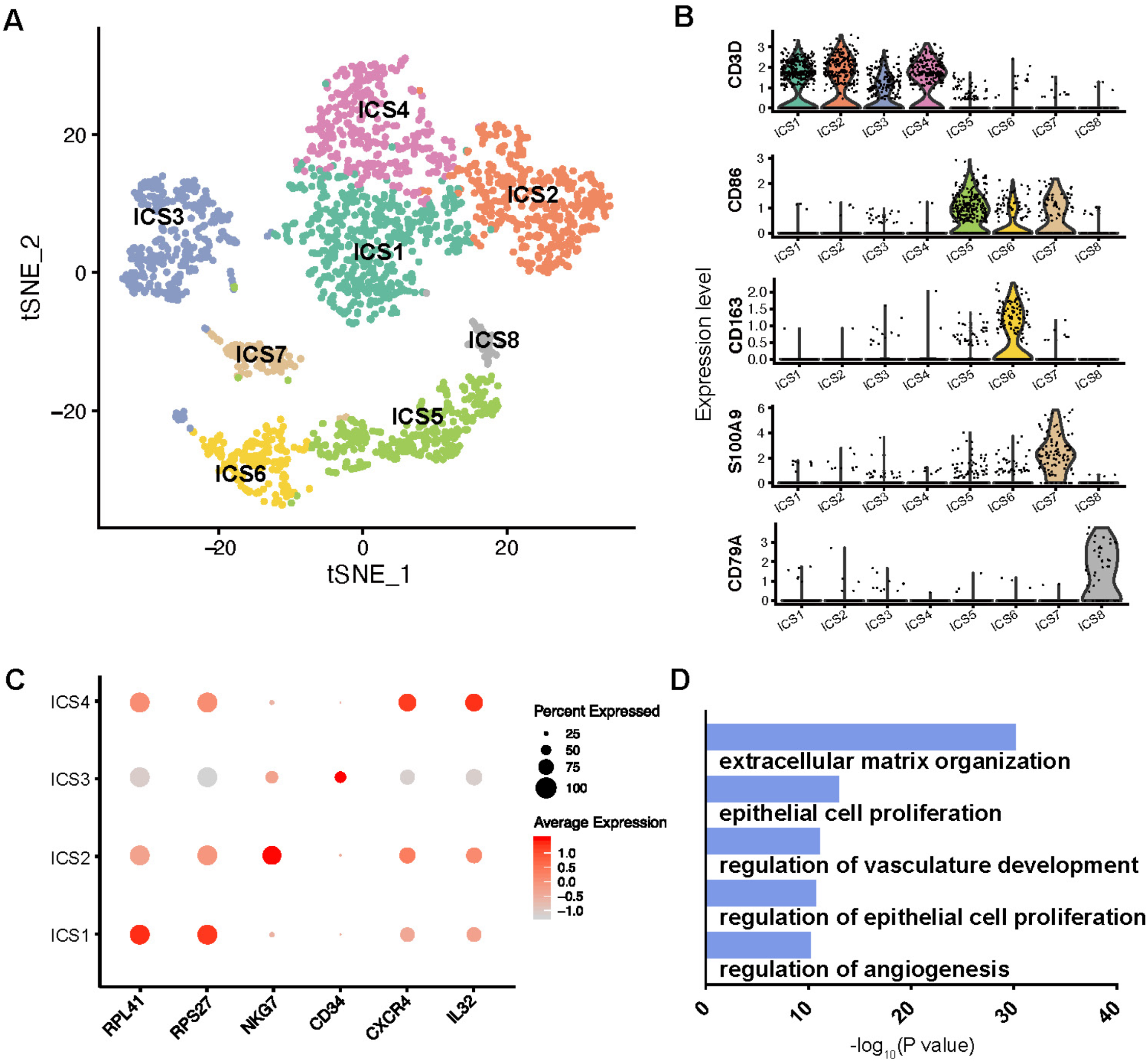
Immune cell heterogeneity of human ASAT. (A) Re-clustering of IC1-1C3 in Figure.1 B identified 8 specific immune cell subpopulations. (B) Violin plots show the expression levels of representative cell-type-specific marker genes across all these 8 immune cell subpopulations. (C) Dot plot of the expression of representative genes across ICS1-ICS4. (D) GO analysis of specific expressed genes of ICS3.

ICS1-ICS4 showed gene expression signatures of T cell and natural killer (NK) cell (Figure. 4B,C). Unsupervised cell type annotation using human primary cell atlas reference (Methods) showed that both of ICS1 and ICS4 were consisted of a mixture of naïve CD8^+^ T cell and memory CD4^+^ T cell (Supplementary Table 3). High expression level of ribosomal protein-related genes distinguished ICS1 from ICS4, while ICS4 expressed higher level of activated T cell related genes *CXCR4* and *IL32* ^20,21^ (Figure. 4C, Supplementary Table 4). ICS2 was annotated as NK cell for the unique marker *NKG7* (Figure. 4C). ICS3 possessed 56% tissue stem cells according to our unsupervised cell annotation (Supplementary Table 3), and highly expressed *CD34* and genes related to ECM organization, epithelial cell proliferation, vasculature development (Figure. 4C,D). Cell quantification demonstrated the most obvious difference between young and old ASAT was the expansion of ICS1 in old tissues (Figure. 5A,B). These results indicate the expansion of a specific T cell subpopulation with active ribosome biogenesis activity during ASAT aging.

**Figure 5.**
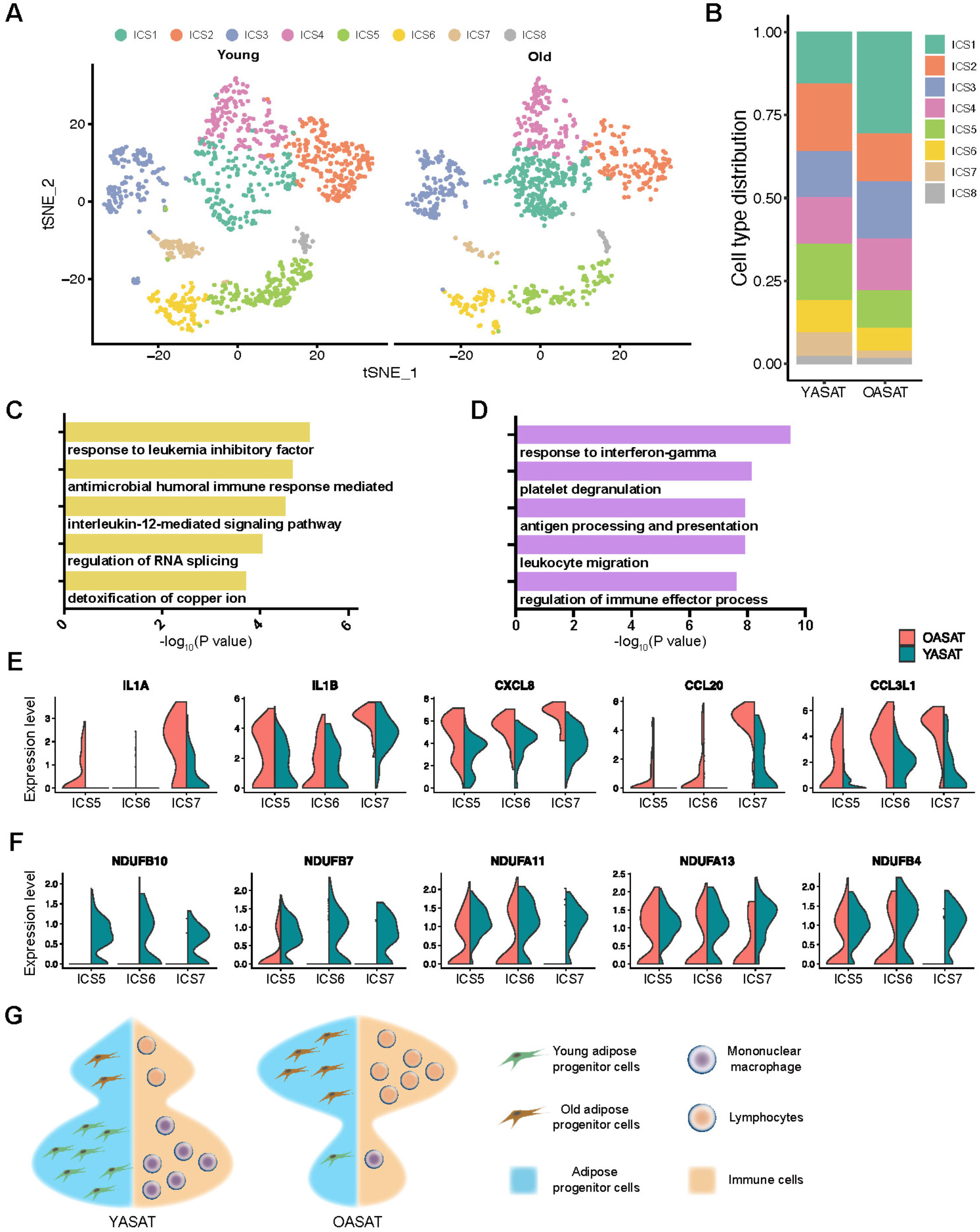
Alterations of immune cell subpopulations during ASAT aging. (A) The t-SNE plots of cell clusters in young and old ASAT split from Figure. 4A. (A) Immune cell type distribution of young and old ASAT. (C) GO analysis of the common upregulated genes in ICS1, ICS2 and ICS4. (D) GO analysis of the common down regulated genes in ICS1, ICS2 and ICS4. (E) Violin plots show the expression levels of chemokine activity-related genes and (F) NADH dehydrogenase-related genes in ICS5-7. YASAT: young abdominal subcutaneous adipose tissue; OASAT: old abdominal subcutaneous adipose tissue. (G) A schematic representation of the changes of immune cell subpopulations in young and old ASAT.

Recent study showed that aging up-regulates specific T cell markers in old VAT ^22^. To investigate whether the expression of these aging-related T cell markers was also upregulated in the expanded ICS1, we analyzed the DEGs between young and old T cell subpopulations. Surprisingly, ICS4 but not ICS1 expressed higher level of aging-related T cell markers *CD44* and *PDCD1* (Supplementary Table 5). Besides, the DEGs analysis revealed that ICS1, ICS2, and ICS4 possessed similar top 10 DEGs during aging (Supplementary Table 5), and in total we found 289 genes were similarly changed by aging in these three subpopulations (Supplementary Table 6). The common upregulated genes were related to antimicrobial humoral response and detoxification of copper ion (Figure. 5C), while genes involved in interferon signaling pathway were downregulated (Figure. 5D). Taken together, these results suggest that T cells in subcutaneous adipose tissue may take a different way responding to aging from visceral adipose tissue, although the cell number was both increased.

ICS5-7 highly expressed *CD86* and antigen presentation-related genes, the markers of monocyte/macrophage. The expression of *CD163* distinguished ICS6 as M2 ^12,23^, and the high expression level of *S100A9* indicated that ICS7 was M1 macrophage ^24^ (Figure. 4B). ICS5 could be identified as dendritic cell according to the unsupervised cell type annotation (Supplementary Table 3). Aging reduced the total proportion of ISC5-7 (Figure. 5A,B) and significantly shifted their transcriptional profile. Genes involved in chemokine activity of ISC5-7 were significantly upregulated upon aging (Figure. 5E), whereas genes related to NADH dehydrogenase was downregulated, suggesting the downregulated oxidative phosphorylation activity, especially in ISC7 (Figure. 5F). These results indicate that aging increases inflammatory level and alters metabolic phenotype of monocyte/macrophage in human ASAT (Figure. 5G).

The smallest cluster ICS8 was designated to B cell for the expression of *CD79A, JCHAIN*, and *MZB1* (Figure. 4B, Supplementary Table 4). Compared to young ASAT, B cells of old ASAT expressed higher level of *IGHA2, IGHG1, IGHG4* and *IGHA1*, which encoded the heavy chain of IgG and IgA (Supplementary Table 5). IgG produced by B cells is known to be associated with insulin resistance in obese mice ^25^. Thus, our data indicate that B cells may contribute to the glucose intolerance and insulin resistance in aged ASAT.

### Aging resets the cell subpopulation interaction pattern in ASAT

Having defined the alterations of progenitor cell and immune cell populations during aging, we utilized iTALK to perform an unbiased ligand-receptor interaction analysis between these populations ^26^.

We inspected the top 20 chemokines and growth factors mediated ligand-receptor interactions. In young ASAT, APC showed strong secretion activity of chemokine CXCL12, an important adipose environmental factor and growth factor CTGF ^27^. CXCL12 acts on the receptors of lymphocyte subpopulations (ICS1-4, and ICS8), whereas CTGF mostly acts within APC subpopulations (Figure. 6A,C). In keeping with our finding that aging increased the inflammatory level in monocyte/macrophage subpopulations, aging dramatically enhanced the secretion of pro-inflammatory factor IL1B by M1 macrophage (ICS7) and the expression of IL1B receptor IL1R1 by APC subpopulations (Figure. 6B). However, the secretion activity of APC was suppressed in aged ASAT (Figure. 6B). During aging, growth factors-mediated interactions were decreased in immune cell subpopulations, and maintained in APC populations (Figure. 6D). Collectively, our analyses demonstrate that the ligand-receptor interaction activity of ICS1 was decreased despite of its increased cell proportion during ASAT aging, and M1 exhibited the most obvious variation in cell interactome, although its cell proportion was slightly decreased during aging.

**Figure 6.**
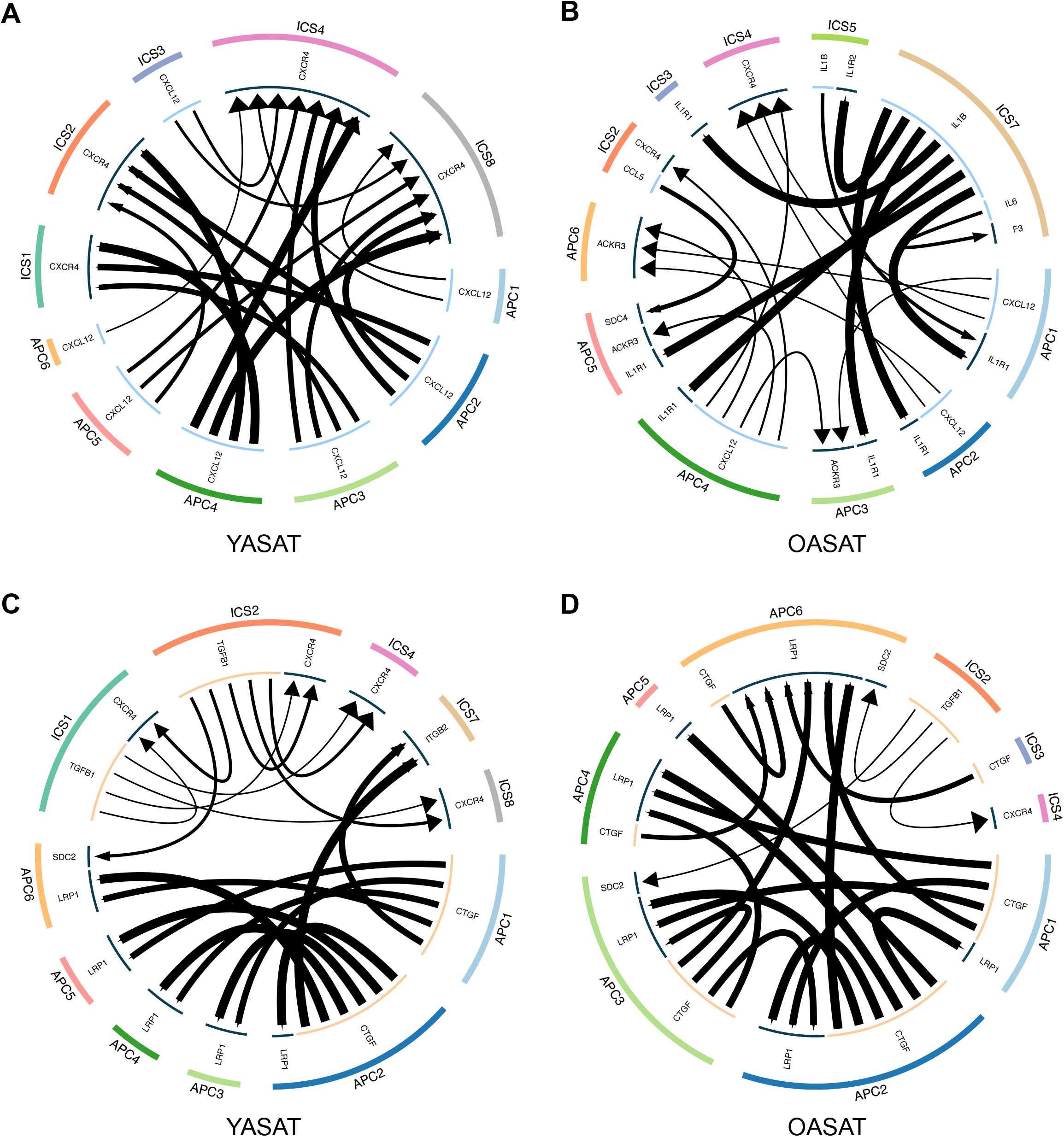
Multi-lineage interactions in young and old ASAT. (A-B) Circus plots showing top 20 chemokines mediated ligand-receptor interaction for all APC and immune cell subpopulations. (C-D) Circus plots showing top 20 growth factors mediated ligand-receptor interaction for all APC and immune cell subpopulations. YASAT: young abdominal subcutaneous adipose tissue; OASAT: old abdominal subcutaneous adipose tissue.

To investigate whether gluteofemoral subcutaneous adipose tissues show the same phenomenon during aging, we performed single cell RNA sequencing on gluteofemoral subcutaneous adipose tissues (GSAT) from young and aged participants. Unsupervised cell clustering and marker-based cell type annotation identified 6 APC populations, 3 IC populations, 1 VEC and 1 SMC population (Supplementary Figure. 4), indicating that overall cell population constitution of GSAT was similar to that of ASAT.

We next asked how GSAT-derived APC respond to aging. Similar to ASAT, aging shifts the distribution of APC populations in GSAT. APC1 mainly composed of cells from old GSAT, and highly expressed cell dysfunctional-related genes (MT2A, MT1E, MT1A and MT1M), which means aging causes the accumulation of a dysfunctional APC subpopulation in GSAT as in ASAT (Supplementary Figure. 5).

Re-clustering of immune cells identified 8 cell subpopulations (Supplementary Figure. 6). Unlike ASAT, the proportion of T cell decreased in aging GSAT, while the proportion of DC cell population was increased. Ligand-receptor interaction analysis showed that APC from YGSAT showed strong secretion activity of chemokine CXCL12 acting on the receptors of lymphocyte subpopulations (ICS1, ICS3, ICS7). In addition, aging dramatically enhanced the secretion of pro-inflammatory factor IL6 by the dysfunctional APC (APC1). These results indicated that APC dominated cell-cell interactions in both young and aged GSAT but mediated by different cytokine (Supplementary Figure. 7).

**Figure 7.**
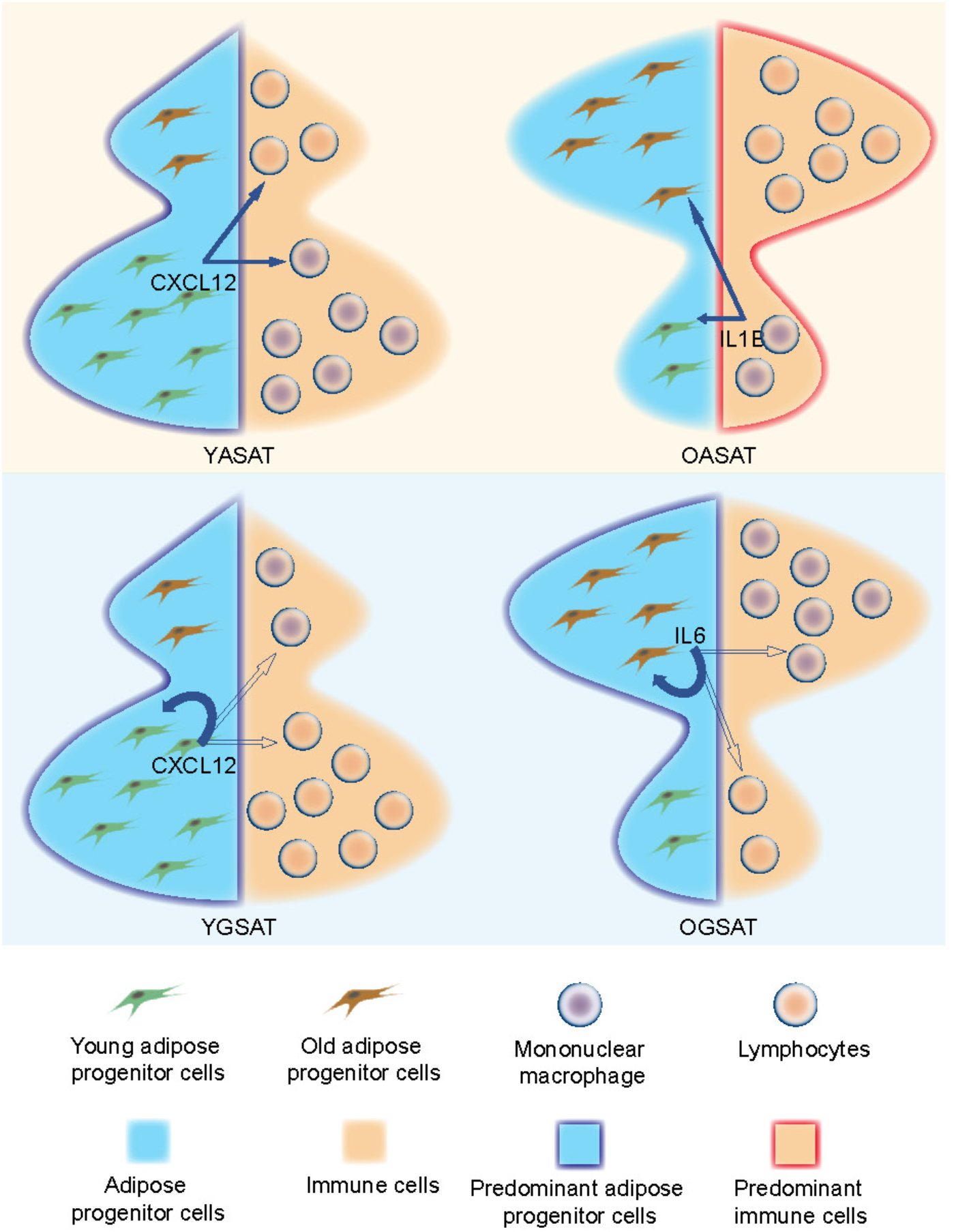
The schematic diagram of cell populations’ crosstalk during the aging process of ASAT and GSAT. In YASAT, adipose progenitor cells dominate the tissue by secreting CXCL12 which acts on immune cells. In OASAT, immune cells dominate the tissue by secreting IL1 which acts on adipose progenitor cells. In YGSAT, adipose progenitor cells dominate the tissue by secreting CXCL12 which acts on both immune cells and itself. In OGSAT, adipose progenitor cells dominate the tissue by secreting IL6 which acts on both immune cells and itself.

### The change of dominating cell population determined the aging of human adipose tissues

In this research, we found aging changes the cell population ratio and crosstalk in adipose tissues (Figure. 7). In both human ASAT and GSAT, aging induced the expansion of dysfunctional aged cells. In young ASAT and GSAT, APC populations dominated the cell-cell interactions through paracrine or autocrine of CXCL12. In old ASAT, immune cell populations secret inflammatory factor IL1 to act on APC populations, indicating an inflammatory tissue microenvironment. In contrast to ASAT, aging didn’t shift the dominating cell population in GSAT. It seems that inflammatory factor IL6 secreted by dysfunctional APC1 of GSAT is the key regulator of the old GSAT tissue microenvironment (Figure. 7).

## Discussion

In this study, we constructed a single-cell aging atlas of human abdominal subcutaneous adipose tissue. The overall cellular component and proportion were similar to a recent published research demonstrating the obese subcutaneous adipose tissue at single-cell level ^28^, which verified the reliability of our data.

Multiple studies have reported the heterogeneity in adipose tissue stem progenitor cells ^14-16^. In addition to the degree of differentiation, there are also different trajectories during stem cell differentiation ^29^. Our results showed the coordinating of maturation stage and differentiation path of APC, and identified at least two trajectories exhibited in ASAT, the ECM secreting APC, and temperature responding APC. Collagen is an important kind of ECM in adipose tissue, it surrounds adipocytes limiting adipocyte hypertrophy and promoting APC adipogenesis ^30^. We found that the ECM-secreting APC (APC4) sharply decreased during aging. The decrease in collagen levels may cause adipose tissue to lose its ability to maintain homeostasis, leaving fat cell hypertrophy in an uncontrolled way, leading to increased hypoxia in adipose tissue and enhanced apoptotic signal, finally suppress adipogenesis of APC. Our data also showed a cluster of stem-like but dysfunctional APC accumulated in elderly adipose tissue, again demonstrating that aging tissue is not always accompanied by stem cells exhaustion, but by arrest of dysfunctional stem cells.

Previous studies demonstrated that the phenotype and function of macrophages are closely related to their metabolic patterns. Pro-inflammatory M1 macrophages mainly rely on glycolysis and present breaks on the tricarboxylic acid cycle. On the contrary, M2 cells are more dependent on oxidative phosphorylation (OXPHOS) ^31,32^. Our data showed that aging enhanced the inflammatory level of adipose tissue macrophages as well as disturbed its OXPHOS process. The synergic effect of inflammation and metabolism deteriorates the elderly macrophages in ASAT.

ASAT aging atlas confirmed the earlier findings that aging is associated with inflammatory macrophages in adipose tissue ^33^. However, our cell subpopulations interaction analysis provided additional support to the idea that M1 macrophages dominated the inflammatory microenvironment of old ASAT. The secretary activity of APC was suppressed by the inflammatory environment during aging, however, the expression level of IL1 receptor responding to the inflammatory signals of M1 was up-regulated. These results also indicate that APC from elderly people may not be suitable for using as stem cell source to treat inflammatory diseases, because it’s susceptible to the inflammatory environment.

In a recently research, the immune functions of structural cells across multiple mice organs were identified ^34^. APC is the structural cells of adipose tissue. Oure data verified the important immune functions of structural cells both in human abdominal and gluteofemoral adipose tissues. Besides, we found that aging could change the immune functions of APC by weakening the strength in abdominal adipose tissue or shifting towards to a proinflammatory status in gluteofemoral adipose tissue. These phenomena suggest that targeting the immune functions of structural cells may be a feasible strategy to treat diseases related to tissue aging.

In conclusion, our study provides a comprehensive characterization of structural cells, immune cells and their crosstalk in human adipose tissues during aging. We emphasize the importance of crosstalk between dominating cell populations and other cell populations in the regulation of tissue microenvironment upon aging. We see our study and large-scale dataset as a starting point and reference atlas for mechanistic explorations in structural cells-immune cells crosstalk mediated tissue homeostasis maintenance.

## Methods

### Study subjects

Human adipose tissues were obtained from patients undergoing a specific surgical procedure with the approval of the Second Affiliated Hospital, Zhejiang University. Specifically, man and woman who had a body mass index between 18.5 kgm^-2^ and 30 kg m^-2^ and required surgery which could harvest abdominal or gluteofemoral subcutaneous adipose tissues were included. For scRNA-seq, the young group included patients between the ages of 16 to 29 years old, the old group included patients between the ages of 68 to 87 years old. The exclusion criteria included pregnancy, liver cirrhosis, inflammatory bowel disease. The characteristics of included patients were in supplementary table 7.

### SVF isolation

To collect SVF cells, adipose tissues were digested at 37°C using collagenase type I (Gibco). SVF cells were collected every 30 mins, and the residual adipose tissues were added to fresh collagenase type I until 95% tissues were digested. Then the SVF cells well treated with RIPA lysis buffer to remove red blood cells. Cells were resuspended with full culture medium (L-DMEM (Gibco)+10% FBS (Gibco)), and 2× freezing medium was added. Finally, cells were placed in programmed cooling box at -80°C for 24 hours, and then stored in liquid nitrogen until analysis.

### scRNA-seq

The scRNA-seq experiment was performed using the Chromium Single Cell 3′ Solution v2 platform (10x Genomics), following the manufacturer’s protocol. SVF cells were thawed. After calculating, the same number of young or old SVF cells were mixed in one tube respectively, and dead cell removal kit (Miltenyi Biotec) was used to improve cell quality. scRNA-seq libraries were prepared using the Chromium Single Cell 3′ Reagent Kit v2. Libraries were sequenced by HiSeq X Ten (Illumina) system.

### Quality control and analysis of single-cell data

The10x Genomics Inc. software package CellRanger (v2.1.0) and the GRCh38 reference genome were used to perform sample de-multiplexing, alignment and quantification of unique molecular identifiers (UMI). The UMI matrices produced by CellRanger were used for downstream analysis by using R (version 3.6.0) and the Seurat package (version 3.1.3) ^35^. Gene expression matrices were filtered to remove cells with >10% mitochondrial genes and < 500 genes. After quality control filtering, data were normalized to remove the effects of the number of genes detected per cell, the number of counts, and the percentage of mitochondrial reads. Normalized data of young subcutaneous adipose tissue were combined into one object and integrated with data of old subcutaneous adipose tissue. Variable genes were discovered using the *SelectIntegrationFeatures* function with nfeatures = 1000. Integration anchors across all samples were discovered using the *FindIntegrationAnchors* function command with default parameters. The *IntegrateData* function was run on the anchor set to integrate all samples with default arguments. Dimensionality reduction was performed with *RunPCA*. Then t-stochastic neighboring embedding method (tSNE) dimensionality reduction was carried out and Shared Nearest Neighbour (SNN) graph constructed using dimensions 1-18 (for ASAT) or 1-15 (for GSAT) as input features and default PCA reduction. Cell clustering was performed on the integrated assay at a resolution of 0.5. SingleR with default parameters was used to generate annotation for each cell cluster ^36^. Differentially expressed genes among clusters was identified by using the function *FindAllMarkers* and examined by Wilcox test, only test genes that are detected in a minimum fraction of 25% cells in either of the two populations. Violin plots, dot plots, and tSNE plots for the given genes and cell populations were generated by using the *VlnPlot, DotPlot*, and *FeaturePlot* functions, respectively.

### Multi-lineage interactome analysis of single-cell data

Cell-cell interaction analysis was performed based on the scRNA-seq data by using iTALK ^26^. Top 50% highly expressed genes were used for further analyses. The software built-in database containing a total of 2,648 unique ligand-receptor interacting pairs were used to identify significant interactions. The top 20 significant interactions were visualized based on the R package circlize ^37^.

### Transcriptional noise analysis

Transcriptional noise between each ASAT cell type derived from old and young donors was quantified based on previous work using the gene expression profiles ^18,38^. Firstly, to remove the effects of the differences in total UMI counts all cells were downsampled to have equivalent number of total counts. Downsampling was also performed on the cell numbers so that equal numbers of young and old ASAT cells in each cell type were used. Then, all the genes were assigned to ten equally sized bins by their mean expression, and the top and bottom 10% of genes were excluded for further analyses. Next, the 10% of genes with the lowest coefficient of variation within each bin were selected. The down-sampled UMI count matrix was reduced to these genes and transformed by square root. Then, the Euclidean distance between each cell and the mean expression level of each cell-type within young and old group was determined. Spearman’s correlation coefficients were calculated on the subsampled UMI count matrix between all pairwise cell comparisons within each cell type and age group. Wilcoxon’s rank sum test was used to evaluate the association between transcriptional noise and age within each cell type. The *p* values were adjusted for multiple testing using the FDR procedure Bonferroni–Hochberg.

## Data availability

The datasets generated during the study are not publicly available for now but will be available before publication.

## Code availability

The code to reproduce the analyses and figures described in this study is available upon reasonable request.

## Acknowledgements

This work was supported by the National Key R&D Program of China (2017YFA0104900), the National Natural Sciences Foundation of China (31830029). We would like to thank The Core Facilities of Zhejiang University-University of Edinburgh Institute for technical assistance, as well as Xuanyuan biotechnology company for their support in nano-indentation.

## Author contributions

W.Z. and J.L. designed and performed experiments and analyzed the data. W.Z. and J.L. wrote the manuscript. X.H. collected human adipose tissues. X.Y. assisted with manuscript writing. H.W.O conceived ideas and oversaw the research program.

