## Supplementary figures for "The shifting of dominating roles between structural cells and immune cells are key regulators of human adipose tissues aging"

### Supplementary figures and legends

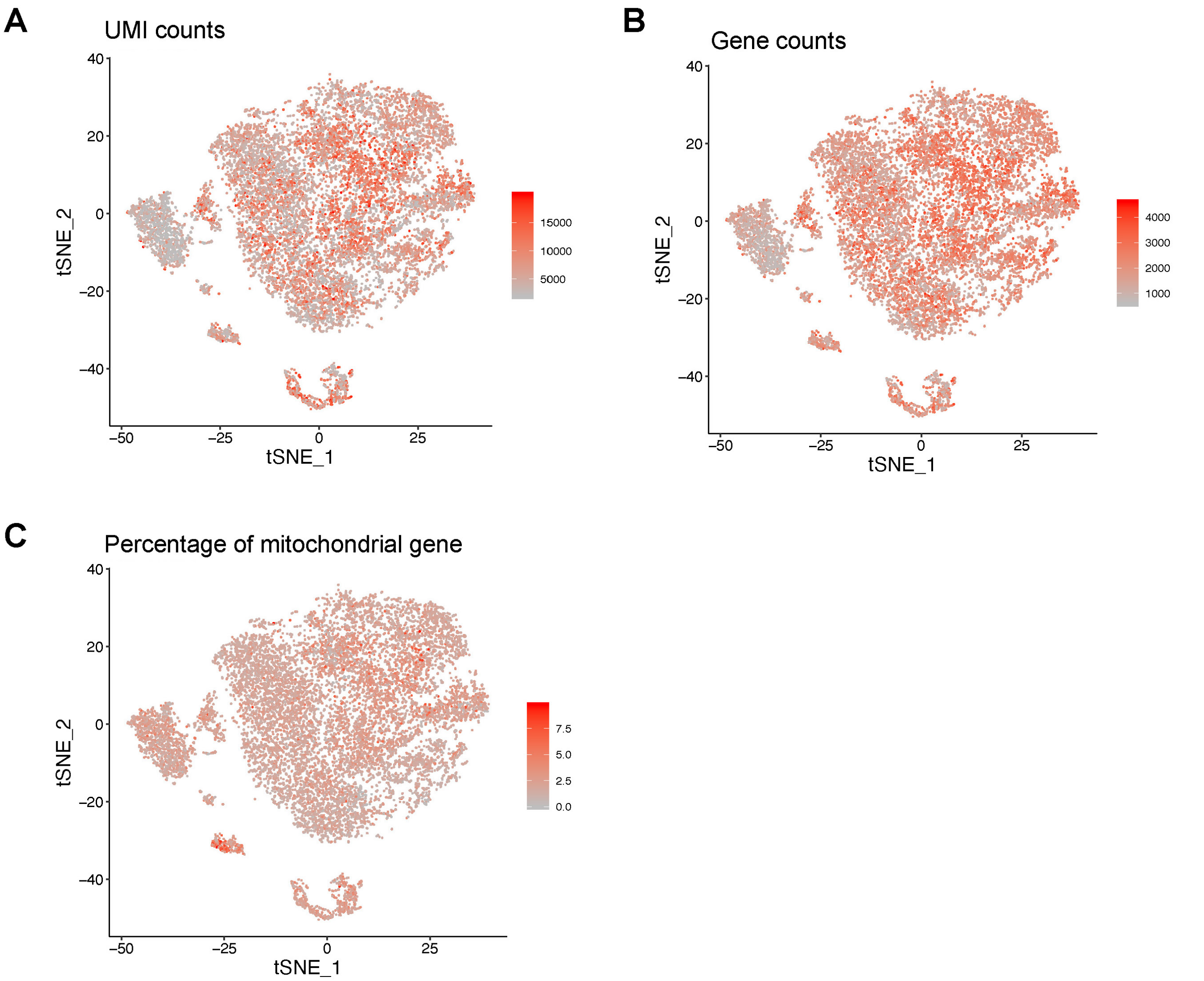


**Supplementary Figure. 1** nUMI, nGene and mitochondrial gene distribution. (A) Cells are colored based on the total number of UMIs. (B) Each cell is colored based on the total number of genes expressed. (C) Cells are colored based on the percentage of mitochondrial gene expression.

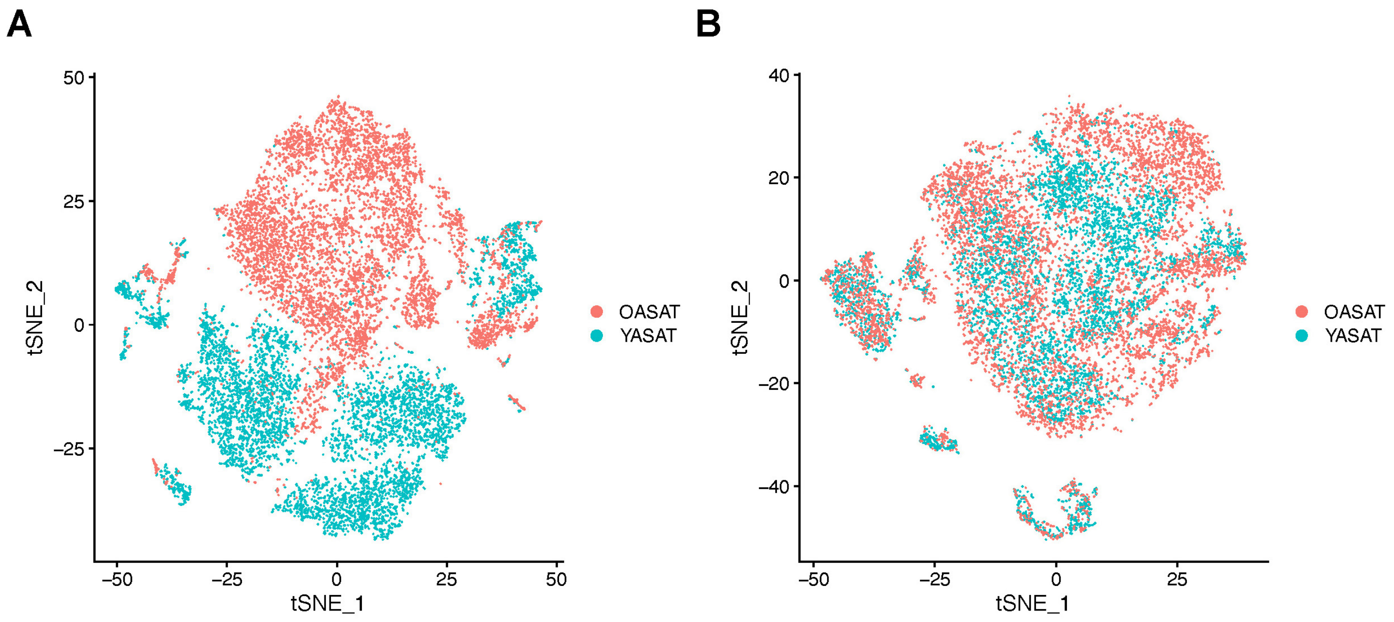


**Supplementary Figure. 2** The t-SNE plots of human ASAT cells split between young and old conditions, before (A) and after (B) alignment.

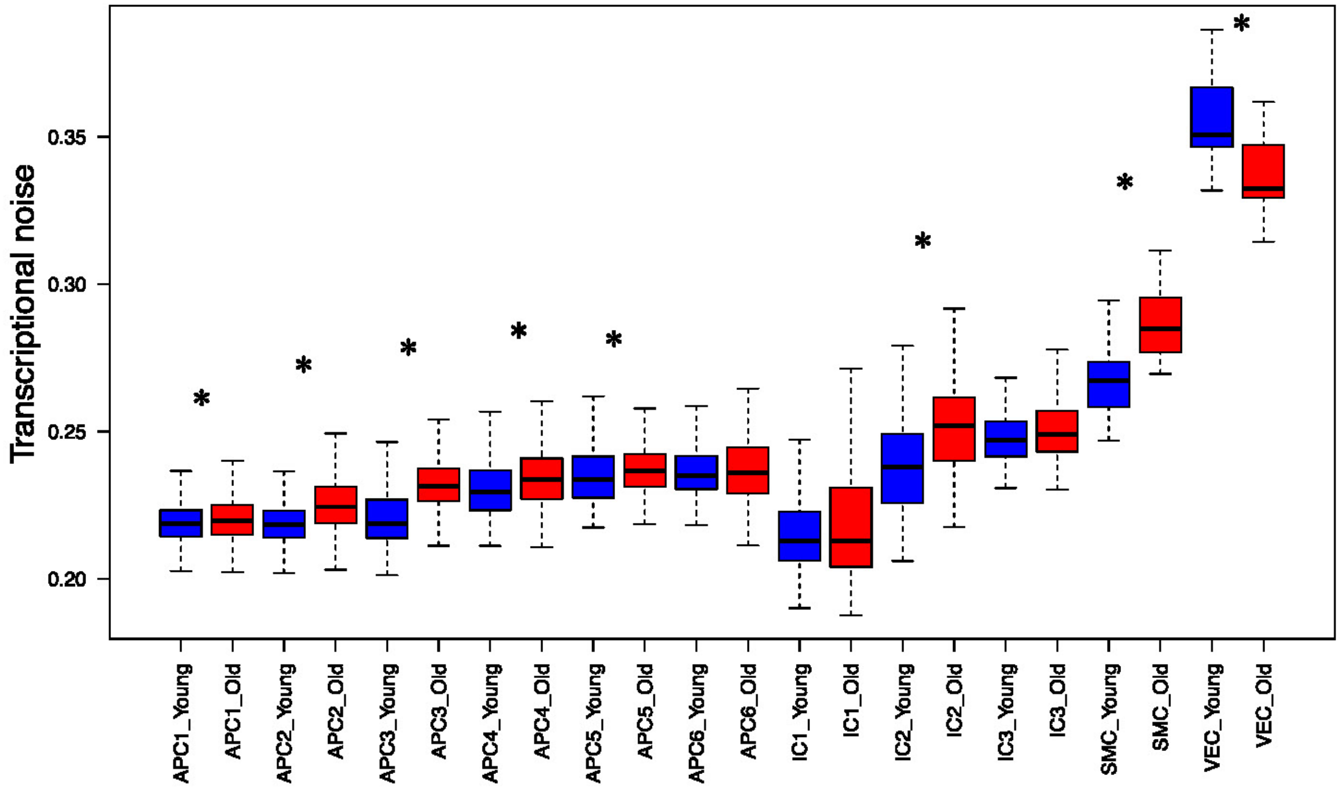


**Supplementary Figure. 3** Transcriptional noise analysis of cell clusters in young and old ASAT. For all boxplots, the box represents the interquartile range, the horizontal line in the box means the median, and the whiskers represent 1.5 times of interquartile range. Asterisk indicates significant changes (assessed by Wilcoxon's rank sum test, adjusted p value < 0.05).

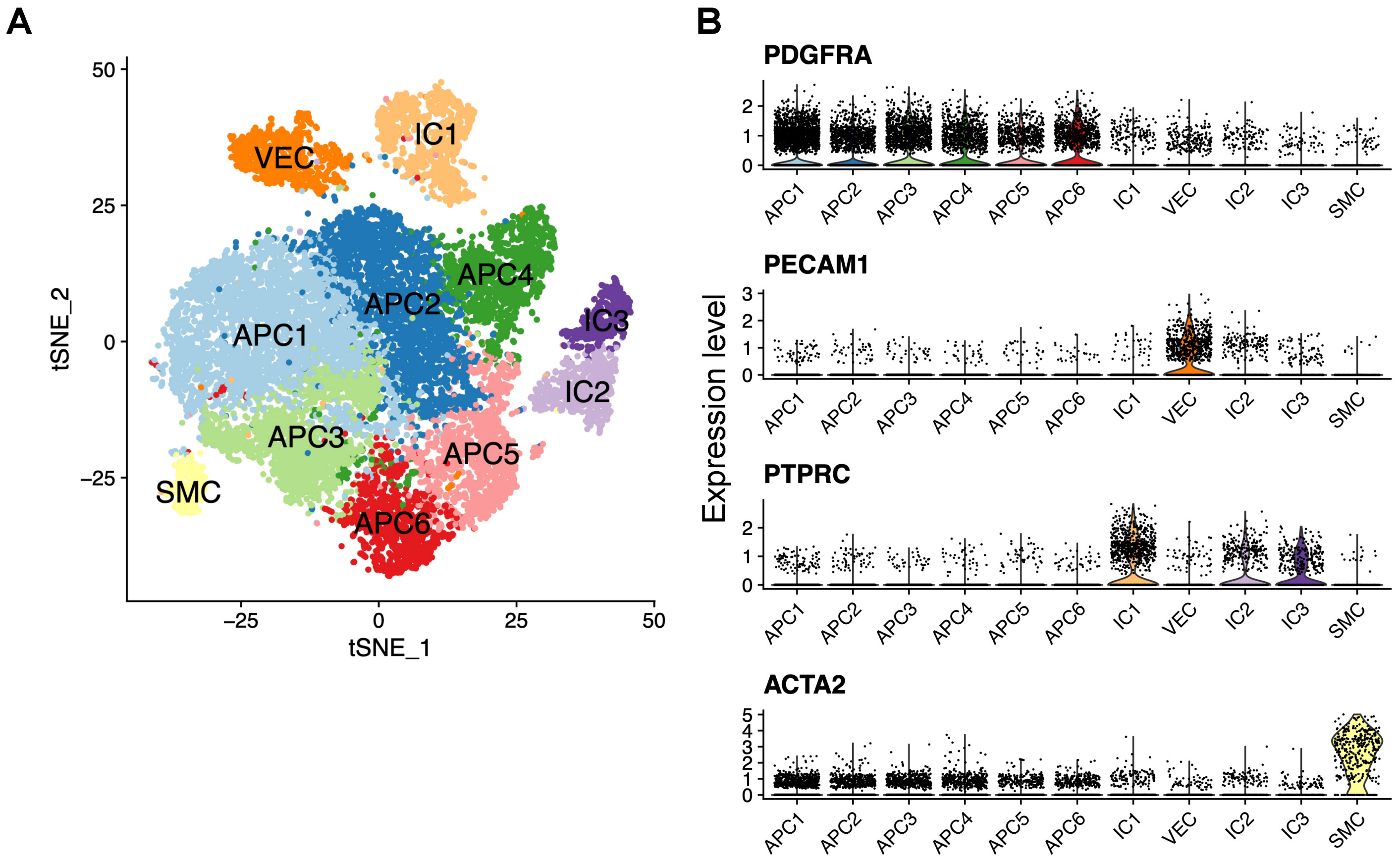


**Supplementary Figure. 4** t-SNE plot of GSAT and marker gene expression. (A) The t-Distributed stochastic neighbor (t-SNE) plot shows unsupervised clustering of GSAT single-cell transcriptomes. (B) Violin plots show the expression levels of representative cell-type-specific marker genes across all 11 cell types.

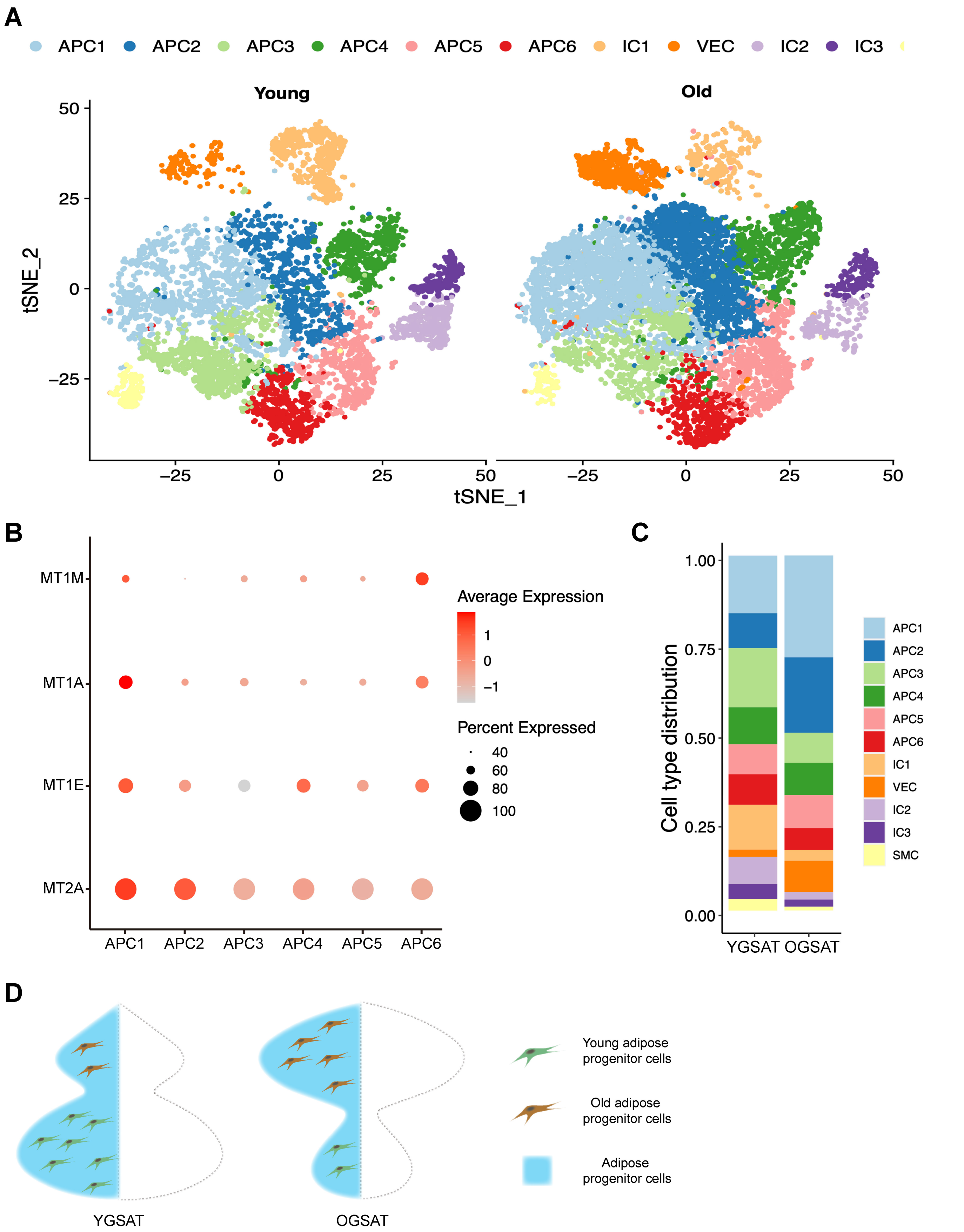


**Supplementary Figure. 5** Alterations of APC subpopulations during GSAT aging. (A) t-SNE plot of cells clusters from Supplementary Figure. 4 split into the cells from young and old GSAT. (B) Dot plot of the expression of metallothionein genes across APC populations. (C) Cell type distribution of young and old GSAT. (D) A schematic representation of the changes of adipose progenitor cell subpopulations in young and old GSAT.

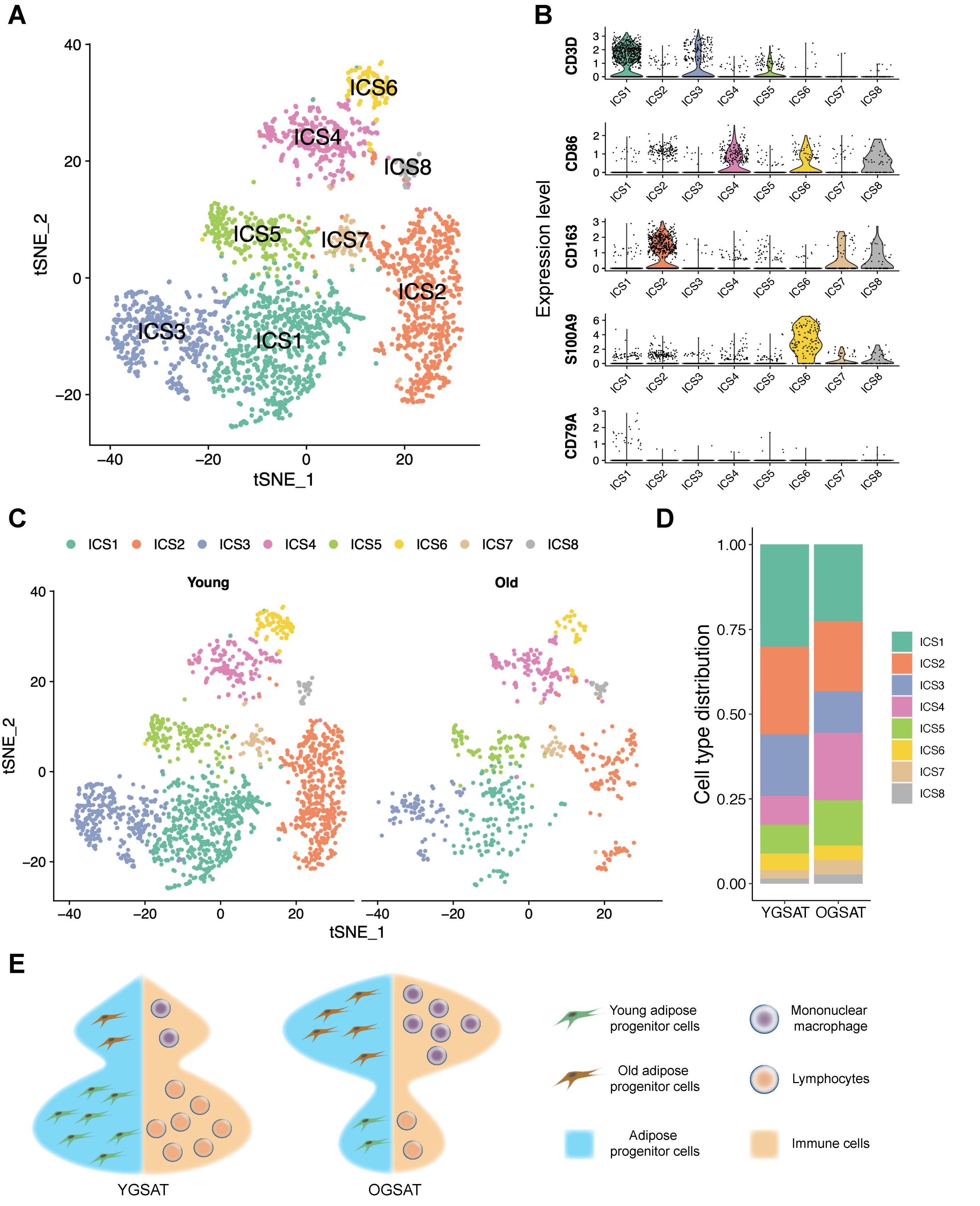


**Supplementary Figure. 6** Alterations of immune cell subpopulations during GSAT aging. (A) Re-clustering of IC1-IC3 in Supplementary Figure. 4A identified 9 specific immune cell subpopulations. (B) Violin plot showing the distribution of expression levels of well-known representative cell-type-enriched marker genes across all these 9 immune cell subpopulations. (C) t-SNE plot split into the cells from young and old GSAT. (D) Immune cell type distribution of young and old GSAT. (E) A schematic representation of the changes of immune cell subpopulations in young and old GSAT.

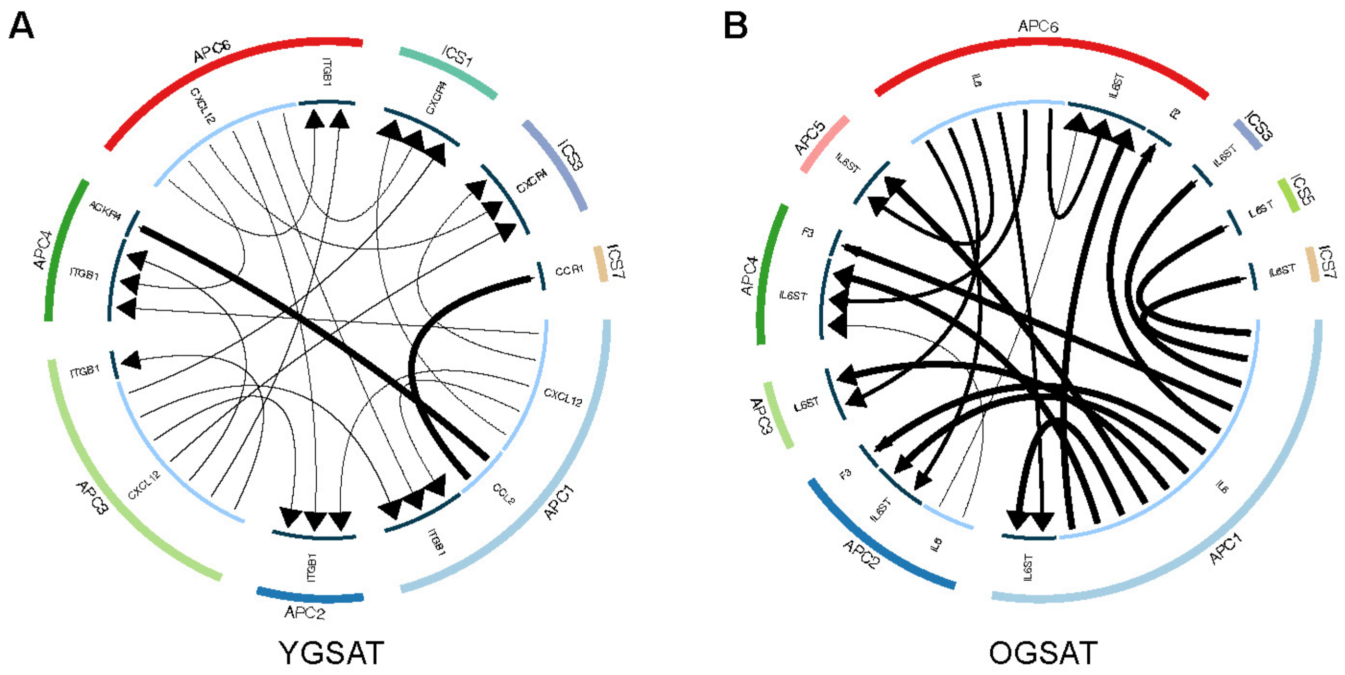


**Supplementary Figure. 7** Circus plots showing top 20 chemokines (A) and growth factors (B) mediated ligand-receptor interaction for all APC and immune cell subpopulations. YGSAT: young gluteofemoral subcutaneous adipose tissue; OGSAT: old gluteofemoral subcutaneous adipose tissue.
